# Camel milk affects serum metabolites by modulating the intestinal microflora

**DOI:** 10.1101/2023.12.18.572112

**Authors:** Haitao Yue, Jiaxue Zhang, Ruiqi Wang, Luyu Zhao, Yuxuan Kou, Runye Li, Zhengyang Yang, Yurong Qian, Xinhui Li, Xiao Wang, Pazilaiti Yasheng, Jieyi Wu, Xiangxiang Xing, Lei Xie, Hao Niu, Gangliang Chen, Jie Yang, Ying Liu, Tian Shi, Feng Gao

## Abstract

Gut microbes play a vital role in human health and are influenced by numerous factors including diet, genetics, and environment. (Fermented) Camel milk, which is abundant in nutrients and lacks allergenic proteins, has been consumed for its edible and medicinal properties for centuries. Research on camel milk’s impact on gut microbiota and host metabolism is still limited. The results found that sour camel milk contained various beneficial bacteria such as *Lactobacillus helveticus, Acinetobacter lwoffii, Eubacterium coprostanoligenes* group, Lachnospiraceae, which could be transported to the recipient’s intestines by diet. This study specified that the transportation of microbiome happened both intra- and inter-species and played a principal role in the formation of progeny gut microflora. An investigation on type 2 diabetic rats revealed that the composition of gut microflora and serum metabolites of those fed with high-dose camel whey was closer to that of the normal. *Eubacterium limnetica*, which can reduce the risk of diseases by producing MtcB protein, was found in the gut microflora of the ones taking camel milk. These results evidenced the high potential of camel milk as a functional food.

## Introduction

Trillions of gut microbes living in the gut play an important role in host biology and diseases(1) . The core human gut microbiota mainly consists of the bacterial phyla Firmicutes, Bacteroidetes, Actinobacteria, Proteobacteria, Fusobacteria, and Verrucomicrobia, of which Firmicutes and Bacteroidetes represent approximately 90% of the gut microbiota community (2). These microbes are closely related to a variety of physiological activities of the human body, including energy transformation, substance metabolism, immune system development, and prevention of pathogen invasion (3).

Shaping of the adult gut microbiome has already been initiated in early life, influenced by factors such as exposure to the maternal microbiome and the mode of delivery, and the early exposure to dietary components (4). Indeed, microbes from food could get gut microbiota regulated. Now, however, numerous of our foods are eaten after heating and cooking, which is nearly asepsis.

A study using apes to decipher the source of gut bacteria proves that most of our gut microbiota have evolved with us for a long time. Moeller found that 2/3 of the major families of gut bacteria in humans and apes could be traced back to their common ancestor about 15 million years ago. When they differentiated from the common ancestor, the gut bacteria split into new strains and coevolved in parallel to adapt to gastrointestinal diseases of different diets, habitats, and hosts (5). Today, these microbes can adapt and help train our immune system, guide the development of our intestines, and even regulate our mood and behavior.

There are many factors that affect the composition and function of gut microbes throughout life, including genetic, sex, age, race, and environmental factors, such as drug use and habitual diet. Among them, diet is the key to modulating abundances of specific bacterial species and their functions. The type, quantity, and balance of nutrients in the diet (carbohydrates, protein and fat, etc.) will affect the composition and number of gut microbes. Conversely, microbes affect the efficiency of food digestion and produce specific metabolites based on dietary substrates, thus affecting the health of other microbes and hosts (6). Therefore, reasonable use of the impact of diet on the gut microbiome is vital for improving human health and the treatment of diseases.

Dairy products are one of the most important protein dietary components in the human diet and part of the official nutritional recommendations of many countries. Consumption of dairy products proved to exhibit positive influences on bone, cardiovascular health, and gastrointestinal microbiome (7). Among various dairy products, camel milk has been consumed for thousands of years for its high nutritional value and health benefits. Camel milk and fermented camel milk production widely spread around Central Asia, Arabian Peninsula, and northwest China (8). Pasteurized camel milk and other derived products, including pasteurized camel milk, ice cream, cheese, camel milk powder, latte, and camel milk soap have been developed and sold in many countries (9).

Camel milk is rich in nutrients (e.g., protein, fat, lactose, vitamins, and minerals) and is a high-potential functional food. Camel milk lacks β-lactoglobulin (β-Lg), which makes it closer to breast milk and less allergic than other animal milk. Camel milk has traditionally been considered to have medicinal properties in some countries and regions (10). For example, camel milk is used to treat jaundice, malaria, and constipation in the Jujiga and Shinile Zones of Eastern Ethiopia and as biomedicine treating several health issues comprising asthma and edema in arid rural regions of Asia and Africa(11).

Besides, camel milk was also known for its anti-diabetic effects. And the low to near zero prevalence of diabetes of the north West Indians who consume camel milk regularly and continuously evidences this (12). Studies proved that camel milk is a beneficial dietary supplement for type 2 diabetes, camel whey protein and its derived hydrolysates and peptides have biological activities against different components of glucose homeostasis, insulin, and glucagon-like peptide-1 (GLP-1) secretion pathways, which can reduce fasting blood glucose, insulin resistance and improve blood lipid levels in diabetes (13). The activation of the AKT pathway downstream of insulin resistance controls the expression and function of glucose transporter 4 (GLUT4) and mediates glucose uptake in principal insulin-sensitive tissues. In addition to that camel milk-derived lactoferrin also acts essentially in glucose transport and uptake, therefore significantly improving insulin sensitivity in type 2 diabetic patients, and exhibits anti-inflammatory and immunomodulatory effects (14).

Fermented dairy is more popular than raw milk because of its advantages of easy digestion, excellent palatability, and easy preservation. The earliest record of the production and consumption of fermented dairy products can be dated back to 5000 BC (15). Until now, fermented dairies such as koumiss and sour camel milk are still popular food products worldwide. Naturally fermented special milk such as camel milk, horse milk, and goat milk is mainly distributed in nomadic settlements (Supplementary Fig.1A). Unlike yogurt products, which are already highly industrialized, the production of naturally fermented dairy remains natural fermentation and handcrafting, which makes a random and diverse microbial taxa. Since the fermented dairy is usually eaten without heating, it largely retains the original microbial community (16). Given that these foods are consumed by such great popularity and are very close to the preindustrial human diets, which inspire us to whether these fermented products act as the vehicle transferring and functioning the microbial community into the recipient’s gut (Supplementary Fig.1B).

35 million camels exist in the world according to data from the Food and Agriculture Organization of the United Nations (FAO) in 2020, in which 89% are *Camelus dromedarius* distributed in North Africa, West Asia, and Australia, the other 11% are *Camelus bactrianus*, mainly distributed in Central Asian countries, including China and Mongolia (8). Though *Camelus bactrianus* is less in quantity, their milk is more abundant in protein, fat, and dry matter content than that of *Camelus dromedarius* (17). China owes rich camel resources, among which Bactrian camels in Xinjiang account for more than 50% of the total. The regulatory effect of camel milk and its functional factors on the body have been widely reported, but few reports on how camel milk affects the host by regulating the gut microbiota or whether as a vector transferring the microbial community into the recipients. Based on second-generation sequencing technology and bioinformatics methods, sampling Bactrian camel milk in Xinjiang, China, this study first scouted the microbial composition and origin of camel milk. Then, the rat model was applied to explore the regulating effect on the gut microbiota of the rats and humans when carrying camel milk intervention. And the microbial floras of camel milk producers and consumers were analyzed to investigate whether microbes could be transferred between species. Finally, the serum metabolism was inspected by LC-MS to investigate the consequence on the host’s metabolism when gut microbiota is regulated by camel milk.

## Results

### Microbial composition and source analysis in camel milk

In this study, the microbial composition of camel milk and sour camel milk from the Darbancheng area (Geographical coordinates 80°57’-88°28’E, 43°22’-43°50’N) and Fuhai County (Geographical coordinates 87°00’-89°04’E, 45°00’-48°10’N)of Xinjiang was firstly analyzed (Fig. 1A). Located on the northern slopes of the Tianshan Mountains, these two cities have been inhabited by nomadic herders since ancient times and have a history of camel breeding and camel milk as food for thousands of years. It was found that Firmicutes and Proteobacteria were the dominant phyla, and the differences between region samples were relatively high. Proteobacteria was dominant in Darbancheng camel milk, while in Fuhai camel milk was Firmicutes. Additionally, the microbes of sour camel milk and camel milk were distinct. One possible hypothesis was that some microbes decrease or even disappear during fermentation, leaving almost all Firmicutes and Proteobacteria. And the microbial composition of fermented milk was also quite different, which was consistent with the distinctness of raw milk from the two regions.

**Fig. 1:**
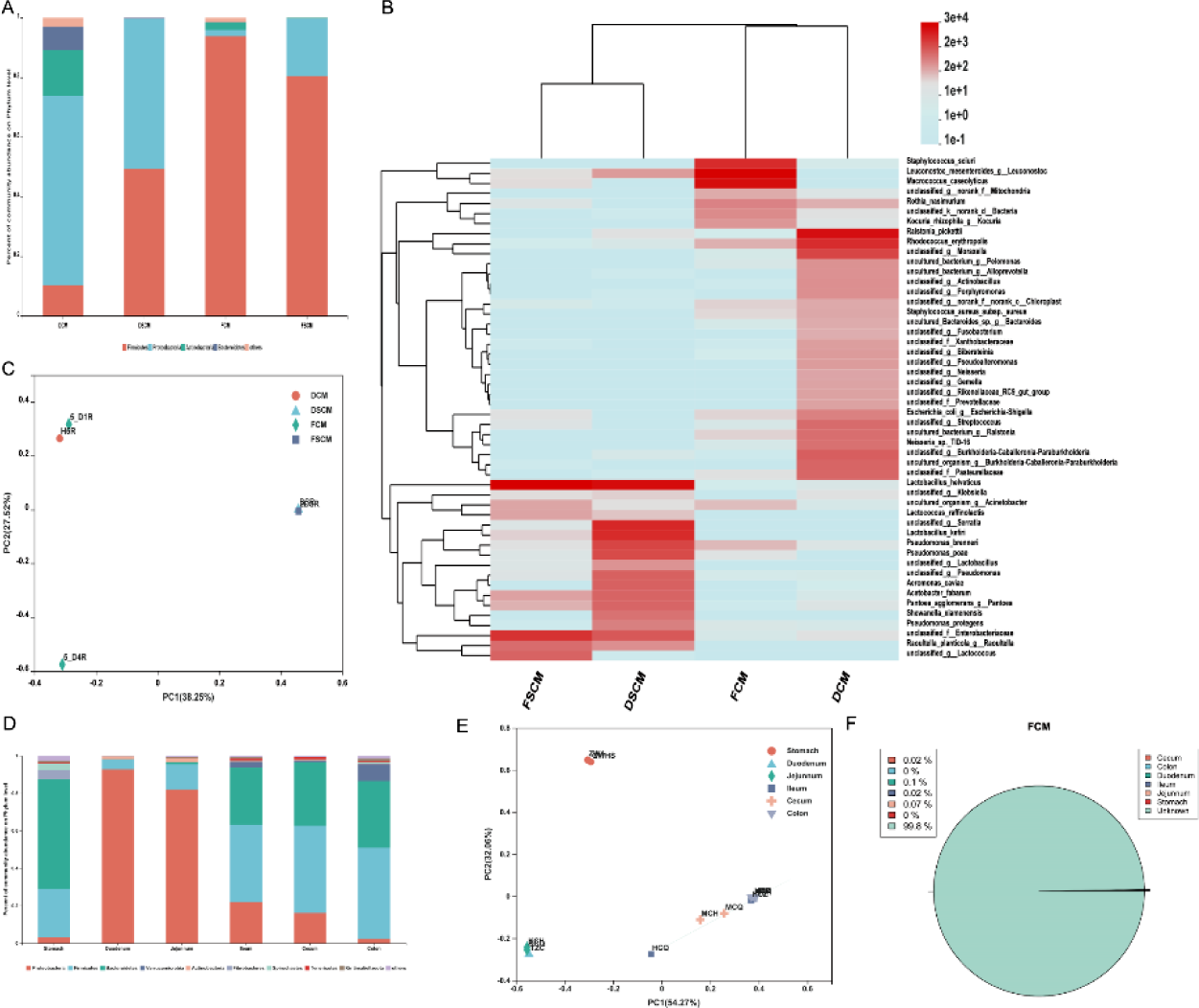
Analysis of microbial diversity in camel milk, sour camel milk, and camel digestive tract. (A) Bar plots of the phylum taxonomic levels in camel milk and sour camel milk. Relative abundance is plotted for each group. (B) Heatmap of microbial communities at the species level in camel milk and sour camel milk. (C) Principal coordinate analysis (PCoA) using Bray-Curtis metric distances of beta diversity in camel milk and sour camel milk. (D) Bar plots of the phylum taxonomic levels in different parts of camel digestive tract. (E) Principal coordinate analysis (PCoA) using Bray-Curtis metric distances of beta diversity in different parts of camel digestive tract. (F) Traceability of camel digestive tract microbes in camel milk.

The dominant bacterial species of camel milk samples from the two regions were different at species level as well (Fig. 1B). Specifically, *Ralstonia pickettii*, *Rhodococcus erythropolis*, and *Moraxella* had the highest abundance of camel milk microbiota in Darbancheng aera, while *Staphylococcus sciuri*, *Leuconostoc mesenteroides,* and *Macrococcus caseolyticus* were dominant in Fuhai County. However, the dominant bacteria of sour camel milk in the two regions were similar to a certain extent, and the most abundant was *Lactobacillus helveticus*. In addition, *Serratia* and *Lactobacillus kefiri* had a high abundance in Darbancheng sour camel milk, and Enterobacteriaceae in Fuhai. The beneficial bacteria in camel milk and sour camel milk were still the majority, such as *L*. *helveticus*, *L*. *kefiri*, and Enterobacteriaceae. The principal coordinate analysis (PCoA) also showed that microbial community structure of camel milk and sour milk was significantly different (Fig. 1C), which was consistent with the result of community composition analysis.

There were many kinds of beneficial bacteria in camel milk and sour camel milk, it’s still unclear whether these bacteria come from the camel’s digestive tract during the lactation process. We analyzed the bacterial composition of the camel’s digestive tract. The results revealed that the microbes in various parts of the camel’s digestive tract were mainly concentrated in Proteobacteria, Firmicutes, and Bacteroidetes (Fig. 1D). Bacteroidetes predominated in stomach, Proteobacteria in duodenum and jejunum, and Firmicutes and Bacteroidetes in ileum, cecum, and colon. PCoA also showed that the camel ileum, cecum, and colon had similar endophytic bacterial community structure, as did the duodenum and jejunum, while the endophytic bacterial community structure of the gastric and intestinal samples were contrasting (Fig. 1E). The results indicated that the microbial composition of different parts of the camel’s digestive tract had certain differences. Compared with the camel milk microbes, the abundance of Bacteroidetes in the camel’s digestive tract was higher, and the microbial composition in camel milk was more similar to that of the duodenum and jejunum. Moreover, only 0.21% of the microbes in camel milk came from the camel digestive tract, mainly from the duodenum and jejunum (Fig. 1F).

### Analysis of the gut microbiota of young camels

Considering that the baby camel is mainly fed on the breastmilk, we hypothesized that their gut microbiota is initially affected by the female camel. Therefore, we compared the fecal microbes of the baby camel and their mothers’. Not surprisingly, their dominant phyla both were Firmicutes, Bacteroidetes, and Actinobacteria. Compared with camel milk microbes, there were more Bacteroidetes and less Proteobacteria, while compared with camel digestive tract endophytes, there were more Actinobacteria and less Proteobacteria (Fig. 2A). The samples from the two regions were also quite different. The dominant phyla in camel feces in Darbancheng were Firmicutes and Bacteroidetes, and the relative abundance of Bacteroidetes in young camels was higher while Firmicutes was lower compared with female camels. In Fuhai camel feces, Firmicutes and Actinobacteria dominated. Compared with female camels, the relative abundance of Actinobacteria was higher while Firmicutes were lower. Therefore, it could be speculated that camel milk had a certain influence on the gut microbiota of young camels.

**Fig. 2:**
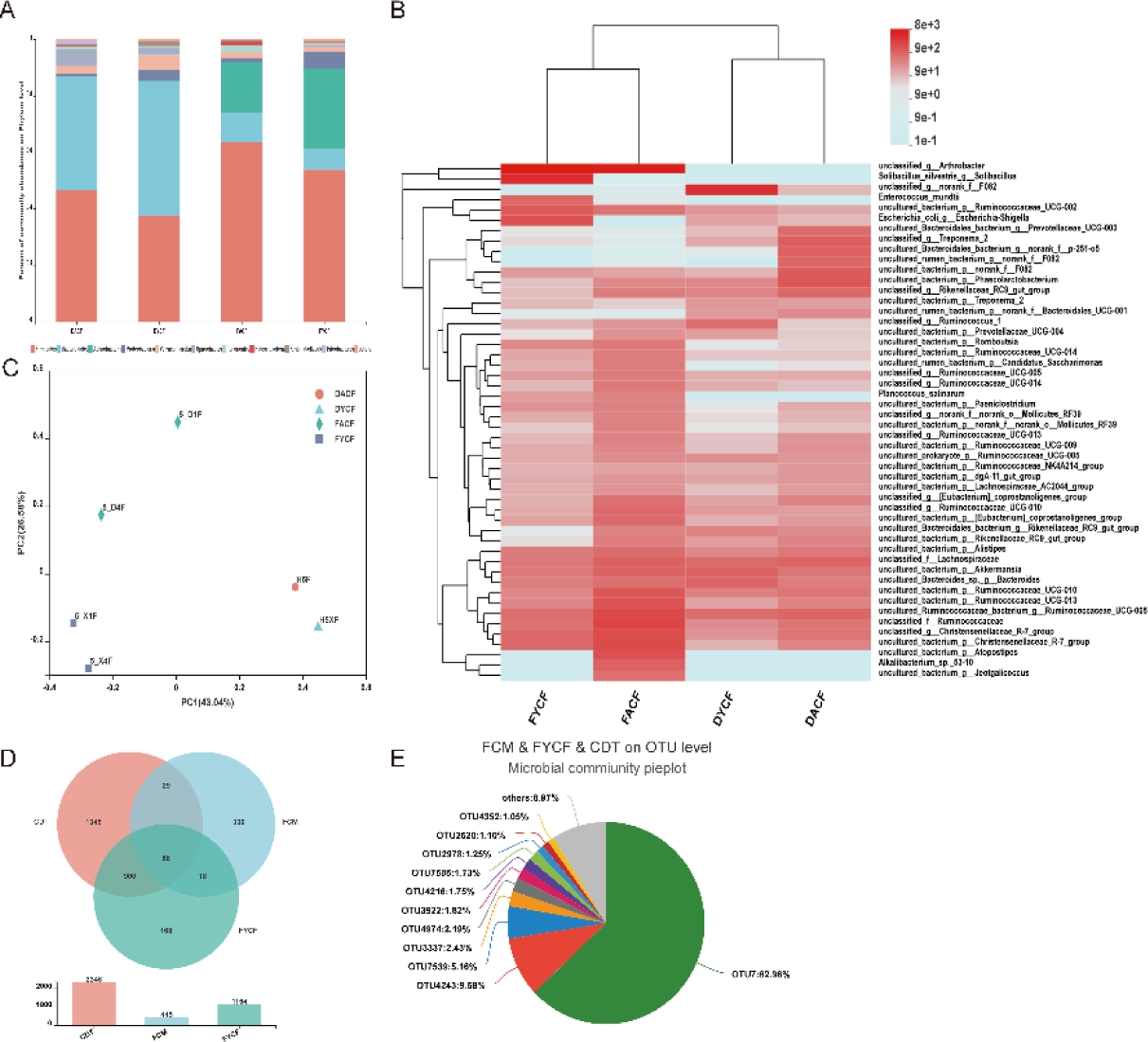
Analysis of microbial diversity in camel feces of female and young camels. (A) Bar plots of the phylum taxonomic levels in camel feces. Relative abundance is plotted for each group. (B) Heatmap of microbial communities at the species level. (C) Principal coordinate analysis (PCoA) using Bray-Curtis metric distances of beta diversity. (D-E) The Venn diagrams of microbes in camel digestive tract, camel milk and young camel feces (D) pie chart of common microbial distribution at the OTU level (E).

At the species level, the microbial composition of the same type of samples from different regions was relatively similar, as well as some differences (Fig. 2B). For example, F082 is the most abundant in camel feces in Darbancheng, while very low in Fuhai. *Arthrobacter* and *Solibacillus* were the most abundant in camel feces in Fuhai, but almost absent in Darbancheng. It was worth noting that *Akkermansia*, with a high abundance in camel feces, was initially separated from human feces and more and more studies had shown its conducive effects on the body. The PCoA analysis revealed that the microbial community structure of camel fecal samples from different regions and the fecal samples of female camels and young camels in the same region both had differences (Fig. 2C), which was consistent with the results of community composition.

The camel fecal microbes shared similarities with camel milk and camel digestive tract, camel milk might be the vector of female camel microbes transferring to the young. 68 OTUs were found in the intersection of the camel digestive tract, camel milk, and young camel fecal microbes at the OTU level (Fig. 2D), which were mainly *Escherichia coli* (OTU7), *Arthrobacter* (OTU4243), Ruminococcaceae UCG-005 (OTU7539), *Bacteroides* (OTU3337), *Akkermansia* (OTU4974), Prevotellaceae UCG-003 (OTU3922), Ruminococcaceae UCG-005 (OTU4216), *Bacteroides* (OTU7505), *Romboutsia* (OTU2978), *Ruminococcus gauvreauii* group (OTU2520), and *Paeniclostridium* (OTU4352) (Fig. 2E). These evidenced that camel milk was a vector transferring microbes from the female camel to their cubs.

### Composition and changes of rat gut microbiota under the regulation of camel milk

Camel milk did modulate the gut microbes of young camels, so did it also affect the gut microbiota of other organisms? We used rats as models to apply camel milk to STZ-induced type 2 diabetic rats to investigate camel milk regulation effect on gut microbiota, along which bovine milk and metformin were used as controls. In the early stage of the experimental group, we compared the effects of camel milk and its components on blood lipid metabolism in diabetic mice, respectively. Investigations unveiled that camel whey performed better hypoglycemic and lipid-lowering activity and liver protection effect than raw milk, skim milk, casein, and camel whey protein (17). Besides, using raw milk was not effective in type 2 diabetic rat model, so we chose camel whey and bovine whey as the diet of type 2 diabetic rats in follow-up experiments.

### Analysis of the composition of gut microbiota in rats

Alpha diversity index detection specified that the Shanno and Chao indexes increased in rats given whey and metformin, proving their effects on improving gut microbiota diversity (Fig. 3A-B). The Shannon and Chao index of the group given high-dose camel whey was significantly different from other groups intimating that high-dose camel whey had an impact on the gut microbes of rats. PCoA also showed that the gut microbiota structure in rats fed with high-dose camel whey was closer to that of normal rats (Fig. 3C). At the phylum level (Fig. 3D), the gut microbes of rats were mainly concentrated in four phyla: Firmicutes, Actinobacteria, Proteobacteria, and Bacteroidetes, in which Firmicutes accounted for the highest proportion with a relative abundance range of 71.0% - 78.63%. Compared with the gut microbiota of diabetic rats, the abundance of Firmicutes and Proteobacteria of rats fed whey decreased, and Actinobacteria and Bacteroidetes increased, but the bovine whey group just exhibited limited variation. While the abundance of Firmicutes in the positive drug group increased, demonstrating that the disparate influences of whey and metformin on the gut microbiota of rats. So were the changes in the family and genus level. Furthermore, beneficial bacteria such as Lachnospiraceae (Fig. 3E) and *Bifidobacterium* (Fig. 3F) were notably more abundant in the gut microbiota of rats fed high-dose camel whey than diabetic ones. We also noticed that the abundance of Lachnospiraceae in the gut of rats fed metformin was significantly lower than that of diabetic rats, which also exhibited the inconsistent impact of high-dose camel whey and metformin on the gut microbiota of rats.

**Fig. 3:**
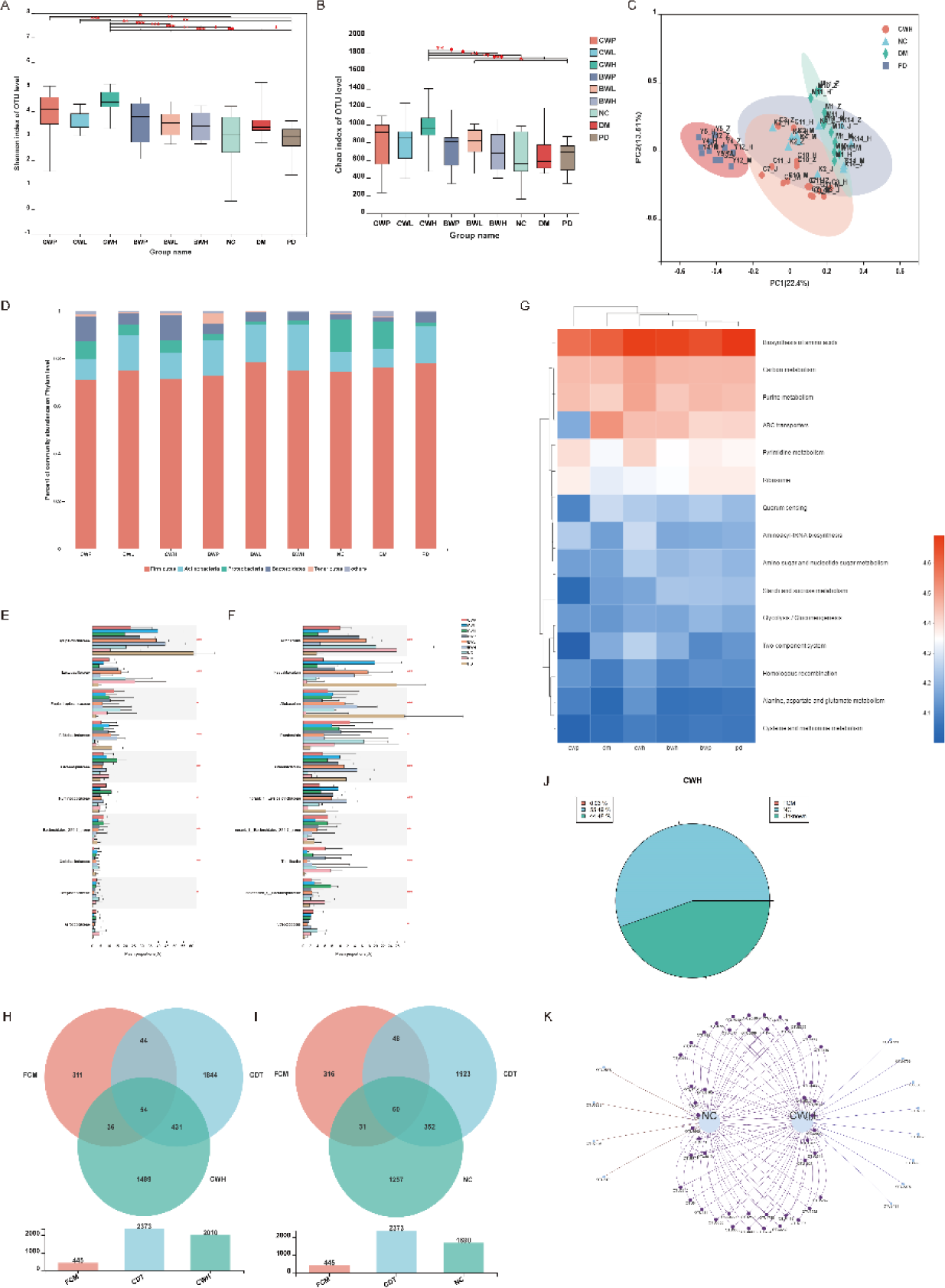
Analysis of gut microbiota in rats. (A-B) Alpha diversity boxplot at the OTU level (A: Shannon index, B: Chao index. * 0.01 < P ≤ 0.05, ** 0.001 < P ≤ 0.01, *** P ≤ 0.001). (C) Principal coordinate analysis (PCoA) using Bray-Curtis metric distances of beta diversity at the OTU level. (D) Bar plots of species abundance of gut microbiota population at the phylum level. (E-F) The significant test of intestinal microbes at the family (E) and genus (F) level (showing the top 10 abundance. * 0.01 < P ≤ 0.05, ** 0.001 < P ≤ 0.01, *** P ≤ 0.001). (G) Heatmap of KEGG Pathway A C B A C B F E G D K I H J Level 3 functional abundance of gut microbes in different groups of rats. (H-I) The Venn diagram of the intersection of OTU levels of camel digestive tract, camel milk and rat gut microbes (H: high dose camel whey group, I: normal control group). (J) The traceability of camel milk microbes in the gut of rats in the camel whey high-dose group and normal control group. (K) The distribution of OTUs at the intersection of camel digestive tract, camel milk and rat intestines.

In addition, different functions of various microbes also affected the metabolic process of the host. The pathways abundance of Biosynthesis of amino acids, Carbon metabolism, Purine metabolism, Pyrimidine metabolism, Aminoacyl-tRNA biosynthesis, Amino sugar, and nucleotide sugar metabolism in the high-dose camel whey group were higher than those in other groups and were comparable to those in the positive drug group (Fig. 3G). Among them, Biosynthesis of amino acids pathway was dominant in all groups. This pathway involved the synthesis of a variety of amino acids, which had a great regulatory effect on glycolipid and energy metabolism. Taking the number of microbes involved in this pathway, 8001 species were noted in the high-dose camel whey group, 3447 in the positive drug group, and only 1467 in the diabetics. It indicated that these microbes might resist the high glucose environment of the host through the synthesis and metabolism of their amino acids, and the effect of high-dose camel milk was more effective than that of metformin. This is consistent with Dekkers’s research, which confirms that metformin treatment is related to the profound changes in intestinal microflora and bacteria carrying genes that can promote amino acid and carbohydrate metabolism (18). The results revealed their abundance became more remarkable when feeding high-dose camel milk and metformin.

### Interspecies transfer of microbes using camel milk as a vector

We proved that camel milk had a positive regulatory effect on the gut microbiota of rats before, but whether camel milk as a vector remained unconfirmed. The analysis found that 54 OTUs intersected among the camel digestive tract, camel milk, and the rats treated with high-dose camel whey (Fig. 3H), while 50 intersected OTU with the normal control was perceived (Fig. 3I) suggesting that camel milk functioned as the carrier by which microbes were transmitted from camel’s digestive tract to rats. And 55.49% of the gut microbes of the rats given high-dose camel whey group were the same as that of normal rats, only 0.03% of the microbes came from camel milk (Fig. 3J). 8 kinds of bacteria were found directly transmitted from camels to rats, namely *Eubacterium coprostanoligenes* group (OTU4729), Mollicutes_RF9 (OTU4769), *Solibacillus* (OTU3386), Ruminococcaceae_UCG-013 (OTU7886), *Oscillibacter* (OTU8523), *Acinetobacter lwoffii* (OTU8995), Mollicutes_RF9 (OTU3378), and Lachnospiraceae (OTU2491) (Fig. 3K), which also confirmed our conjecture.

### Effects of camel milk-regulated gut microbiota on metabolism in rats

High-dose camel whey exploited a good enrichment and regulation effect on the gut microbiota of rats, this inspired us whether the effect directly relied on synchronizing host metabolism. Therefore, we analyzed the serum metabolites, blood sugar, and body weight of the high-dose camel whey group, diabetes model group, positive drug group, and normal control group by metabolomics, respectively. PCoA analysis (Fig. 4A-B) divulged the significant differences in the results applying positive and negative ion modes of each group. In particular, the groups of rats fed with high-dose camel whey and metformin were nearly those of normal rats but significantly different from those of diabetic rats, indicating the positive ramifications on the metabolism of diabetic rats of both camel whey and metformin.

**Fig. 4:**
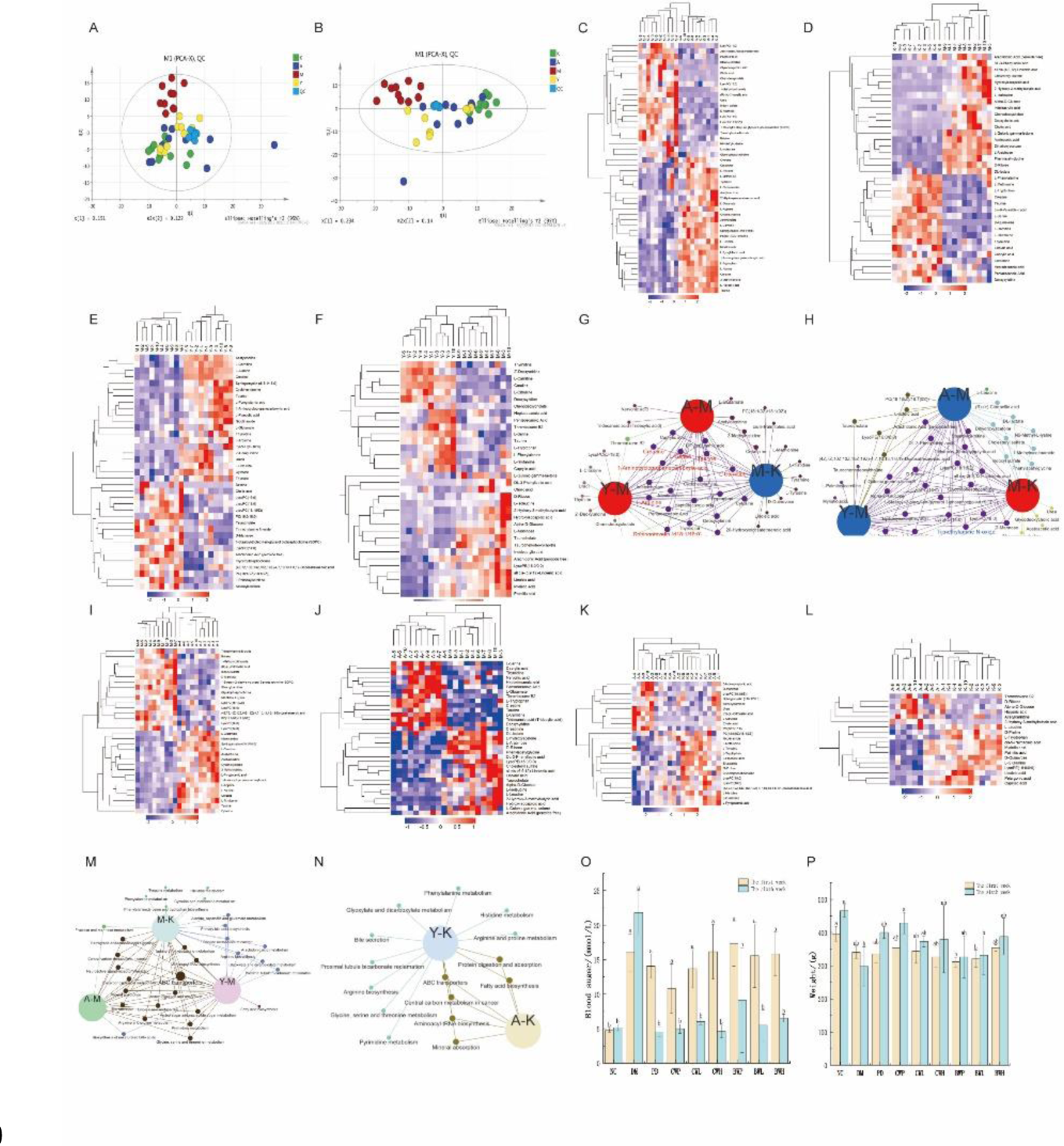
Serum metabolomics analysis of different groups of rats. (A-B) PCA score map of serum samples under positive (A) and negative (B) ion mode. (C-D) Cluster analysis of differential metabolites in serum samples among A-M group under positive (C) and negative (D) ion mode. (EF) Cluster analysis of differential metabolites in serum samples among Y-M group under positive (E) and negative (F) ion mode. (G) Network diagram of up-regulated metabolites in rats fed highdose camel whey and metformin with down-regulated metabolites in diabetic rats. (H) Network diagram of down-regulated metabolites in rats fed high-dose camel whey and metformin with upregulated metabolites in diabetic rats. (I-J) Cluster analysis of differential metabolites in serum samples among A-Y group under positive (I) and negative (J) ion mode. (K-L) Cluster analysis of differential metabolites in serum samples among A-K group under positive (K) and negative (L) ion mode.(M) KEGG pathway network of M-K, Y-M and A-M. (N) KEGG pathway network of Y-K and A-K. (O-P) Blood glucose (O) and body weight (P) of rats before and after six weeks.

Clustering of differential metabolites between different groups showed that the dysregulation of metabolites caused by diabetes was virtually reversed after feeding high-doses camel whey or metformin (Fig. 4C-F, Supplementary Figure). A comprehensive picture of positive and negative ion models indicated that 22 up-regulated metabolites in rats fed high-dose camel whey were the same as those down-regulated in the diabetics, except Creatinine (0.05< p <0.1), the remaining were significantly different (p<0.05); the similar occasion found in the group treated with metformin, in which 24 metabolites up-regulated and all were included in the down-regulates of the diabetics except L-Tryptophan and Pentadecanoic acid (0.05 < p < 0.1) (Fig. 4G). Meanwhile, 33 and 28 down-regulated metabolites in rats fed with high-dose camel whey or metformin were the same as the up-regulated metabolites in diabetic rats, respectively. Other down-regulates were also detected, including Arachidonic acid (peroxide free), DL-3-Phenyllactic acid, LysoPC (14:0), LysoPC(16:0), and LysoPC(18:1(9Z)) (0.05< p <0.1) in camel whey group and Cholic acid, D-Ribose, L-Isoleucine, and Trimethylamine N-oxide in metformin, the remaining were significantly different (Fig. 4H).

Additionally, the number of differential metabolites of the rats fed high-dose camel whey or metformin was significantly reduced, indicating their similar effects on serum metabolites in diabetic rats (Fig. 4I-J). Interestingly, when compared with normal rats, the number of differential metabolites fed high-dose camel whey was less than that of the metformin group. We clustered 4 samples with the normal control group and the results indicated that the metabolites were verged on normal rats, which was consistent with the principal component analysis (Fig. 4K-L, Supplementary Fig. 3).

Furthermore, the KEGG signaling pathways of differential metabolites enrichment in diabetic rats and the rats fed with high-doses camel whey or metformin were mainly concentrated in ABC transporters, Protein digestion and absorption, Central carbon metabolism in cancer, Aminoacyl-tRNA biosynthesis, Mineral absorption and so on. Among them, the pathway of ABC transporters had the largest number of enriched metabolites (Fig. 4M). In this pathway, the transport of monosaccharides, phosphates, and amino acids were affected most by diabetes. Our investigation indicated that feeding high-dose camel whey or metformin could both regulate the transport of monosaccharides (e.g., D-Mannose, D-Ribose, L-Arabinose), phosphate (e.g., L-Serine, L-Alanine, L-Arginine, L-Glutamate, L-Glutamate, L-Isoleucine, L-Leucine, L-Phenylalanine, and Taurine), and amino acid, thereby restoring the effect of diabetes to this pathway (Supplementary Figure 2). Moreover, compared with the rates fed with metformin, the signaling pathways of differential metabolites enrichment between rats fed high-dose camel whey and normal rats was far less (Fig. 4N). And the rats fed with high-dose camel whey performed a similar hypoglycemic rate and weight gain rate to that fed metformin, which was close to the normal control (Fig. 4O-P). Previous reports stated that bacterial functional modules related to amino acid metabolism exhibited stronger positive enrichment in metformin-related species (19). Combining, though with approximative hypoglycemic effect, high-dose camel whey performed extraordinary regulation and restoration on the gut microbiota and serum metabolism of rats than metformin.

### The structure and changes of gut microbiota in people taking camel milk

Previously, we proved that camel milk could affect the host metabolism by regulating the gut microbes of rats. But what would the situation be when it comes to humans remained unexplored. The same method was applied to analyze the human gut microbiota, which could be explored by analyzing the fecal samples, of pastoral herders drinking camel or bovine milk.

The PCoA illustrated that the structure of the gut microbiota of camel milk drinkers was significantly different from those who drank bovine milk or short-time-intakes of camel milk (Fig. 5B). The comparison analysis indicated that the abundance of Firmicute and Bacteroidetes decreased, and Actinobacteria and Proteobacteria increased in the gut microbiota of individuals drinking camel milk compared with bovine milk drinkers (Fig. 5A). What’s remarkable was that even a short-time intake of camel milk performed changes to the gut microbiota. Compared with individuals who experienced a long time of drinking camel milk, the abundance of Firmicute was reduced, and that of Actinobacteria and Proteobacteria increased considerably, reaching comparable. It might be due to the structure of gut microbiota having changed quite and being in an unstable state in the early intervention of camel milk. And at the genus level, *Prevotella*, *Clostridiu*, *Faecalibacterium*, and *Lachnospiraceae* were enriched in the gut microbiota of camel milk-drinking individuals compared with bovine milk-drinkers (Fig. 5C). The main driving force of global differences in intestinal flora seems to come from lifestyle rather than geographical location (20). Pastoral herders adopting a non-industrial lifestyle usually have a more diverse microbiome. The diversity of intestinal flora of non-industrial people appeared earlier and was influenced by the local environment (19).

**Fig. 5:**
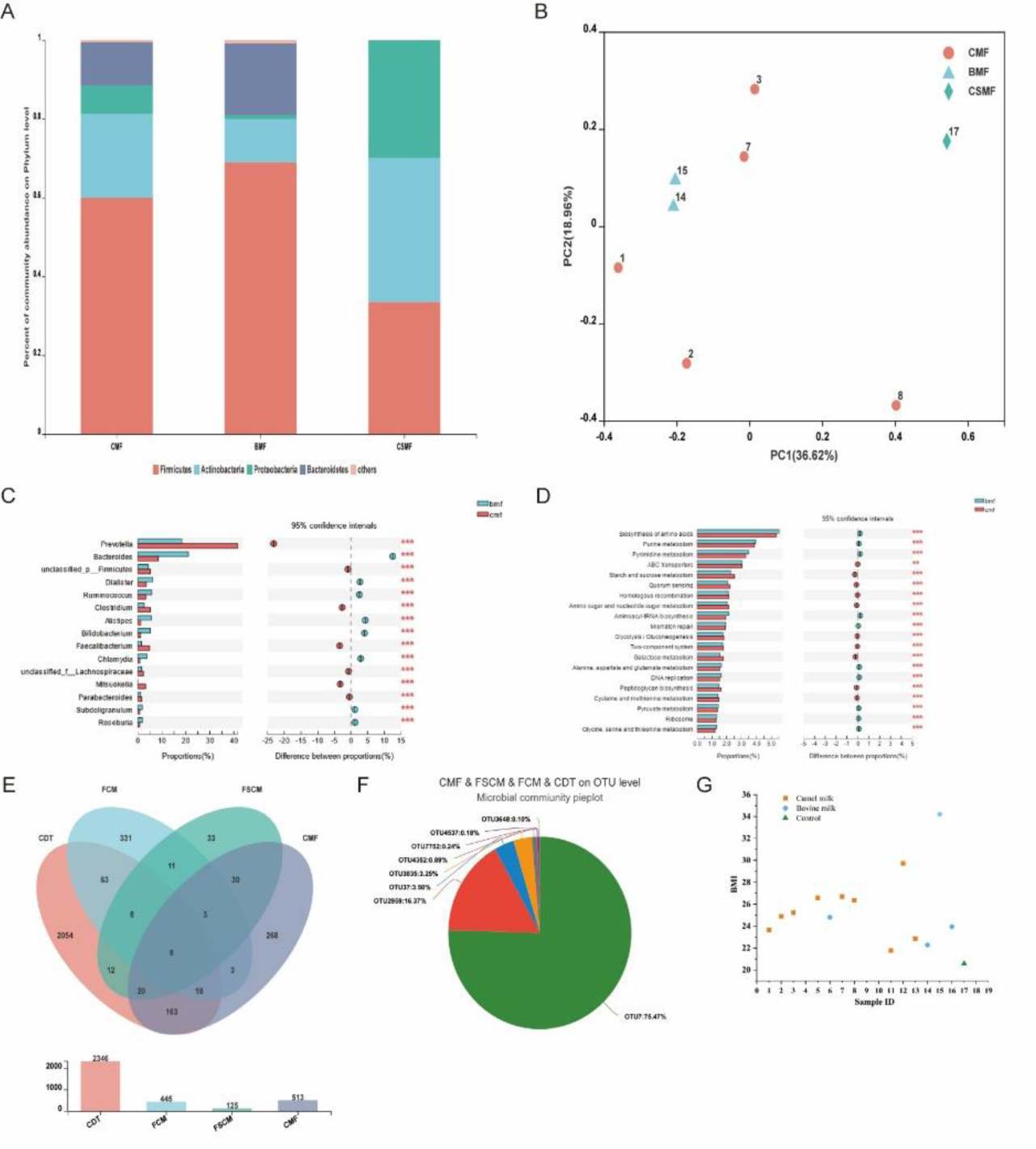
Analysis of the composition and function of human gut microbiota. (A) Bar plots of the phylum taxonomic levels in gut microbes. Relative abundance is plotted for each group. (B) Principal coordinate analysis (PCoA) using Bray-Curtis metric distances of beta diversity. (C) Comparision table of differences in gut microbiota between individuals drinking camel milk and bovine milk at the genus level. (D) Comparision table of differences in microbial KEGG Pathway Level 3 function in gut microbiota of people drinking camel milk and bovine milk. (E-F) The Venn diagrams of microbes in gut microbes who drank camel milk, camel milk and and yogurt (E) and pie chart of common microbial distribution at the OTU level (F). BMI of human (G).

Under the KEGG Level 3 Pathway of the gut microbiota of individuals drinking different milk (Fig. 5D), the abundance of Starch and sucrose metabolism, Amino sugar and nucleotide sugar metabolism of individuals drinking camel milk was higher than that of those drinking bovine milk, while the Biosynthesis of amino acids, Purine metabolism, and Pyrimidine metabolism was the opposite. This was different from the diabetic rats fed with camel whey or bovine whey, possibly caused by the samples being from non-diabetics, and no single dietary intervention was performed. However, it was severe to execute a single dietary intervention on humans, and various aspects could interfere with the state of their gut microbiota. An impressive appearance was that the BMIs of all adult herdsmen were less than 30 except for one, and no exhibited any sign of diabetes (Fig. 5G). Taking the rats’ results combined, it could be concluded that camel milk had a positive effect on the host by regulating the gut microbiota.

Considering hygiene and taste, people usually do not drink camel raw milk directly. Among diverse camel milk products, naturally fermented milk was the closest to raw milk. Therefore, we tracked the microbes transmitted from camel to human through sour camel milk. The exploration indicated that 8 OTUs in the camel’s digestive tract intersected with camel milk, sour camel milk, and human feces (Fig. 5E), including *Escherichia coli* (OTU7), *Prevotella*_9 (OTU37), *Lactobacillus helveticus* (OTU2959), Lachnospiraceae (OTU3648), Enterobacteriaceae (OTU3835), *Paeniclostridium* (OTU4352), *Chloroplast* (OTU4537), and Lachnospiraceae (OTU7752) (Fig. 5F).It has been reported that environment-dependent colonization may play a key role in the transmission of microbiome among individuals, the inheritance of microbiome from parents to offspring, and the coevolution of host-microbiome (21).

### Analysis of the endophytic flora of camel edible desert plants

Unlike cattle, horses, sheep, and other centrally housed animals, camels are well adapted to foraging in the wild, and this traditional feeding practice is retained today in northern Xinjiang, China. Camels feed on a wide range of wild Gobi Desert plants, so we collected 15 species included in the camels’ recipe and analyzed their endophytic bacteria, of which Chenopodiaceae accounted for 60% (Supplementary Table 6). Most of their endophytes were found to be concentrated in Proteobacteria, Firmicutes, and Actinobacteria (Fig. 6A). Among them, Firmicutes was the dominant bacteria in Chenopodiaceae and Zygophyllaceae, Actinobacteria in Amaranthaceae, and Proteobacteria in Asteraceae, Brassicaceae, and Leguminosae. However, even within the same family of plants, the differences in the endophytic community existed in different species (Fig. 6B).

**Fig. 6:**
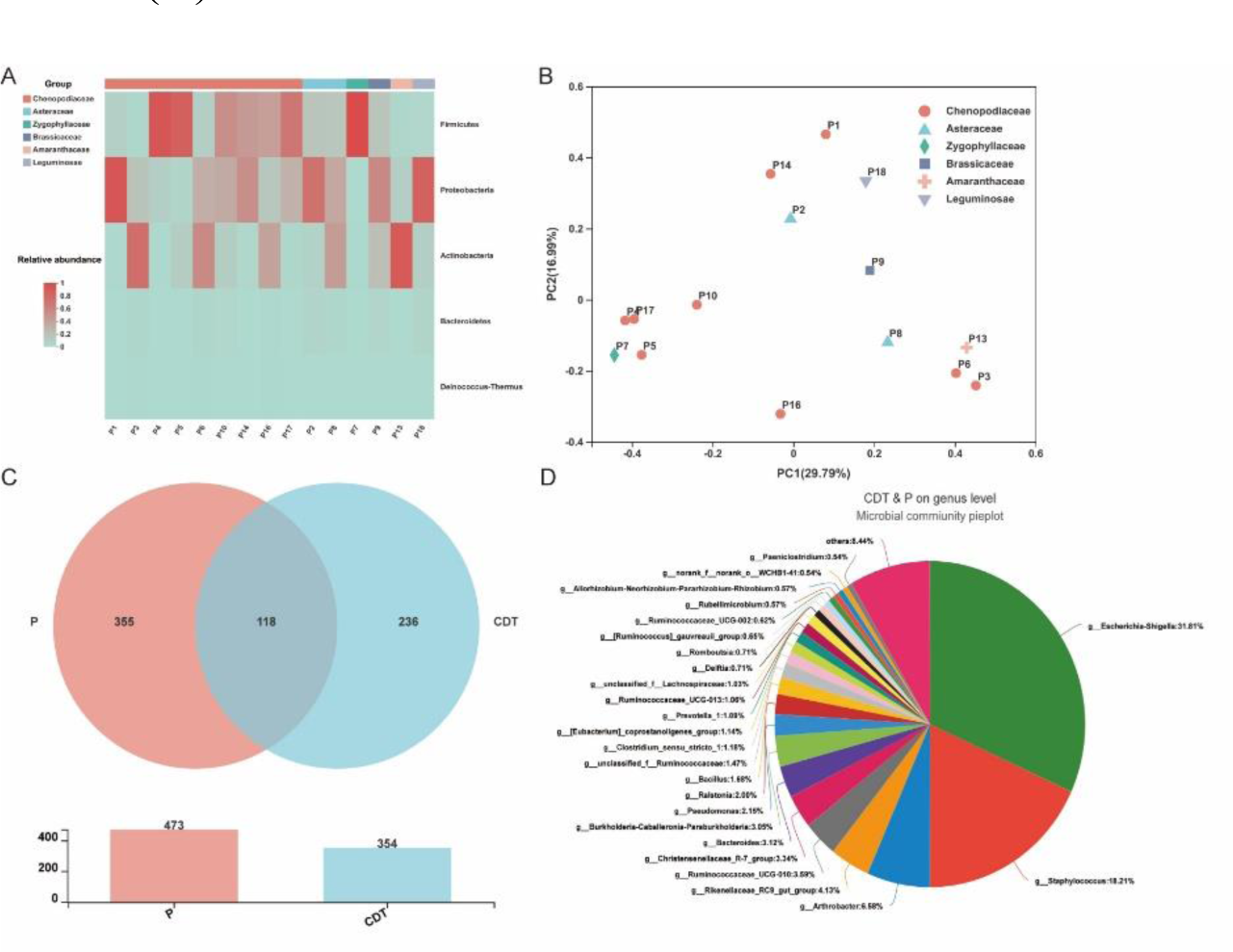
Analysis of the endophytic flora of camel edible desert plants. (A) Heatmap of selected most differentially abundant features at the phylum level. (B) Principal coordinate analysis (PCoA) using Bray-Curtis metric distances of beta diversity. (C-D) The Venn diagrams of microbes in desert plants and camel digestive tract (C) and pie chart of common microbial distribution at the genus level (D).

Microbes can be transmitted using camel milk as a vector in both intraspecific and interspecific. But whether the microbes in the camel’s digestive tract were sourced from their food, the Gobi Desert plants. We compared the microbial genera of these plants and camel digestive tract and found 118 genera at their intersection (Fig. 6C), comprising some with high abundance like *Escherichia*-*Shigella* (31.81%), *Staphylococcus* (18.21%), *Arthrobacter* (6.58%), Rikenellaceae_RC9_gut_group (4.13%)), Ruminococcaceae_UCG-010 (3.59%), Christensenellaceae_R-7_group (3.34%), *Bacteroides* (3.12%), and *Burkholderia*- *Caballeronia*-*Paraburkholderia* (3.05%) (Fig. 6D).

The transmitted microbes found in the above were all within the range of these 118 intersecting genera, and they had variable effects on the physiological and biochemical activities of the body. Previous studies have also indicated that transmission of the gut microbiota can occur between humans and mammalian when they come into close contact, such as predator-prey interactions (22). In short, we speculated that camel milk can act as a vehicle to deliver natural microflora to colonize and function in the gut of the recipient.

## Discussion

In line with our research, we speculated that the camel-rearing way was the prominent drive of the diversity of microbial composition of (sour)camel milk or camel feces. In the Darbancheng area, camels usually have a relatively simple diet because they are mainly raised in captivity and fed with silage based on alfalfa. However, camels in Fuhai are mainly raised by grazing and with a complex diet compromising various natural desert plants.

Camel milk and fermented camel milk are in abundance of nutrients and probiotics. *L. helveticus* existed in sour camel milk samples from both regions, could regulate the structure of gut microbes and promote the host’s health. *L. kefiri*, which had good probiotic potential and was isolated from camel milk in Kazakhstan, could adjust gut microbiota, fight cancer, and inhibit harmful bacteria and alleviate Toxins, and so on(23–26). A beneficial bacterium *Akkermansia* was also found in camel feces. *A. muciniphila* in this genus was a new member of the symbiotic microbiota discovered in recent years, which was of great significance for the prevention and treatment of diabetes, obesity, and cancer(27, 28). Bae et al., reported the identification of a lipid from *A. muciniphila’*s cell membrane that recapitulates its immunomodulatory activity in cell-based assays (29).

Diet possessed enrichment and regulation effects on gut microbiota. When diabetic rats were fed with whey, the diversity of gut microbiota increased similar to that fed with metformin, the relative abundances of Firmicutes and Proteobacteria decreased, and Actinobacteria and Bacteroidetes augmented. Interestingly, the composition of gut microbes of rats fed high-dose camel whey was closer to that of normal rats Compared with bovine whey and metformin, and beneficial bacteria such as *Bifidobacterium* and Lachnospiraceae were more enriched. *Bifidobacterium*, the most reported beneficial genus in type 2 diabetes research, was inversely associated with type 2 diabetes and could be naturally present in the human gut or introduced as a probiotic(30). Hannah C. et al. found that the overall diversity of gut microbiota in people who ate fermented foods increased significantly, in which Lachnospiraceae, Ruminococcaceae, and Streptococcaceae in Firmicutes were the principal, and it was positively correlated with the quantity of consumed (31). This was predominantly consistent with our research but only the increased abundance of Lachnospiraceae and Ruminococcaceae was observed in rats fed camel whey.

We also noticed the functional abundances in the Biosynthesis of amino acids pathway of intestinal flora of rats fed with high-dose camel whey and metformin were higher than that of in other groups. Its synthetic branched-chain amino acids (BCAAs) such as valine, leucine, and isoleucine were associated with insulin resistance (32). The analysis of serum metabolomics found that a large number of amino acids and organic acids were significantly increased in rats fed with high-dose camel whey and metformin, and it had been reported that they had a positive effect on insulin resistance, diabetes, and other metabolic diseases. For example, L-arginine can improve vascular dysfunction in diabetic rats by inhibiting ET-1/Nox4 signaling pathway-related endothelial cell apoptosis (33). L-alanine had a relieving effect on alloxan-induced diabetes, can restore tissue antioxidant properties, and improved kidney and liver damage (34). L-serine was positively correlated with insulin secretion and sensitivity, and continuous supplementation reduced diabetes incidence and insulitis scores, improved glucose tolerance, decreased insulin resistance index, and lowered blood glucose levels in non-obese diabetic rats(35). Taurine is a non-protein amino acid that improves fasting and postprandial blood glucose, serum insulin levels, insulin resistance, β-cell function, and insulin sensitivity(36). It can improve sciatic nerve axonal injury in diabetic rats by activating PI3K/Akt/mTOR signaling pathway (37). Creatine is a nitrogenous organic acid that, when combined with exercise, positively affects glucose metabolism, increases insulin secretion, alters osmolarity, and improves glycemic control by increasing glucose uptake by increasing GLUT-4(38–40). In addition, L-aminocyclopropanecarboxylic acid can also inhibit the memory decline of young mice, enhance the object recognition memory and cognitive flexibility dependent on the prefrontal cortex, and also have a positive regulatory effect on the nervous system (41).

Furthermore, L-carnitine, an amino acid-like amino acid that promoted the conversion of fat to energy, was upregulated in the serum of rats fed high-dose camel whey and metformin. High-dose L-carnitine could increase the total antioxidant status, superoxide dismutase and glutathione peroxidase levels in the pancreas and serum of STZ-induced diabetic rats, and reduce the antioxidant capacity of the rats (42). And substances such as L-carnitine and choline can be metabolized by intestinal flora to produce Trimethylamine (TMA). In the liver, TMA can be converted into Trimethylamine N-Oxide (TMAO) under the oxidation of Flavin containing monooxygenase 3 (Fmo3). TMAO is a vital risk factor for the onset of cardiovascular and cerebrovascular diseases such as atherosclerosis and thrombosis. The level of TMAO in cardiovascular patients is considerably higher than that in healthy people, which also indicates a higher risk of disease(43). Krzycki et al. found that the MtcB protein in *E. limosum* and *E. callanderi* could interact with L-carnitine in the gut, cutting off the methyl group of L-carnitine, thereby preventing L-carnitine from generating TMA and reducing the risk of disease. This study indicated that TMAO levels were significantly down-regulated in rats fed high-dose camel whey, and *E. limosum* was found in the gut microbiota of humans taking camel milk. And the TMA-producer *Anaerococcus hydrogenalis, Clostridium asparagiforme, Clostridium hathewayi, Clostridium sporagenes, Escherichia fergusonii, Proteus penneri, Providencia rettgeri*, and *Edwardsiella tarda* were not found in both rat and human guts fed high-dose camel whey and metformin. Moreover, Sphingomyelin, upregulated in rats fed high-doses of camel whey and metformin, is also an important signaling molecule in eukaryotes. Tofte et al. found that in type 1 diabetes, higher levels of sphingomyelin and specific alky lacyl phosphatidylcholines were associated with lower risk of end-stage renal disease and all-cause mortality (44). A research team from BGI found that genes involved in polysaccharide degradation and sphingolipid metabolism were abundant in *Bifidobacterium* in a functional analysis of 1520 human culturable intestinal bacteria reference genomes. And *Bacteroidetes* also contained a considerable number of genes related to the synthesis of sphingolipids and steroid hormones(45). We also noticed that both genera were highly enriched in the gut microbiota of rats fed high-dose camel whey, and were also presented in the humans.

Our investigation confirmed the vector role of camel milk, by which microbes were transferred into the recipient’s gut and work, including many beneficial bacteria. *Ruminococcaceae*_UCG-005 and *Ruminococcaceae*_UCG- 013 are the dominant genera in the digestive tract of ruminants. They can not only maintain a healthy and stable level of the intestinal tract but participate in the digestion and absorption of remaining nutrients and prevent the loss of nutrients(46). *Bacteroides* and *Akkermansia*, as *Bifidobacterium* mentioned above, are beneficial bacteria genera negatively associated with type 2 diabetes and are worthwhile in glucose metabolism in humans and experimental animals. *Romboutsia* is a butyrate-producing bacteria, and butyrate can inhibit inflammation and regulate intestinal immune function by inhibiting the activity of NF-kB (47). The Harbin Medical University team found that the *E. coprostanoligenes* group in mouse feces mediated the hypolipidemic effect of high-fat diet through sphingosine, an upstream substance in the glycosphingolipid biosynthesis pathway (48). *Oscillibacter*, a stress-sensitive microbial taxa, whose abundance was significantly reduced in patients with major depression. *Acinetobacter lwoffii* is an environmental bacterium isolated from cattle farms known to prevent childhood asthma (49). *Lactobacillus helveticus* is a probiotic. Lachnospiraceae and Enterococcaceae are also beneficial flora of the gut. Gut microbiota, especially Lachnospiraceae and Enterococcaceae, was proven can protect mice against radiation-induced damage to the hematopoietic and intestinal systems, and thus survive lethal doses of radiation. And these beneficial microbes were significantly higher in the feces of leukemia patients with mild radiotherapy side effects (50).

Probiotics are widely used in the prevention and treatment of human diseases or health disorders by rebalancing the host’s gut microbiota. The nine probiotics currently available for baby food include four *Bifidobacterium* and five *Lactobacillus*, which are present in camel milk, sour camel milk, and the gut microbes of rats and humans (51, 52). In addition, prebiotics can selectively promote the metabolism and proliferation of beneficial bacteria in the body, thereby improving the health of the host. An example is that short-chain fatty acids (SCFAs) produced by microbial fermentation of dietary fiber in the gut play important roles in intestinal homeostasis, adipose tissue, and liver substrate metabolism and function, and can prevent type 2 diabetes (T2DM) and Non-alcoholicfatty liver disease (NAFLD) (53).

It is currently accepted that probiotics and prebiotics are provided through suitable food as a vector matrix. Milk protein is a source of both amino acids and health-positive bioactive peptides. Dairy products, in particular, are also major probiotic vectors(54). Compared with bovine milk, camel milk contains lower saturated fatty acids and higher unsaturated fatty acids, showing higher antioxidant capacity and angiotensin-converting enzyme inhibitory potential when simulating gastrointestinal digestion(9). And camel whey protein shows higher protection against gastrointestinal diseases than bovine whey (55). Our results confirmed camel whey’s “prebiotic-like” effect for enriching and regulating gut microbes and as a microbe-transferring vector, which had positive effects on gut microbiota and body health. Our recent separation work revealed that four proteins (e.g., lactoferrin, α-lactalbumin) included in camel whey were responsible for its unique effect. However, the purification and quantitative investigation are limited by the low productivity of camel milk. An alternative approach to obtain these proteins is heterologous production using microbial chassis, which can be further exploited for quantitative study. We also have explored the heterologous production of the camel milk-sourced functional proteins and the details will be elaborated in future publications.

## Materials and methods

### Sample collection and processing

**Desert plants**: collected from pastoral area 635, Fuhai County, Altay Region, Xinjiang, China (Geographical coordinates 80°57’-88°28’E, 43°22’-43°50’N). Herdsmen grazed twice a day, usually at 9:00 am and 4:00 pm, that observed and tracked the types and quantities of plants the camels feed in the desert, and then collected the stem and leaf tissues of the plants and put them in sterile bags. See Supplementary Table 6 for sample information.

**Camel digestive tract**: collected from healthy camels in Fuhai County, Altay, Xinjiang, China. See Supplementary Table 7 for sample information.

**Camel milk, camel feces, and sour camel milk**: collected from Darbancheng (Geographical coordinates 87°00’-89°04’E, 45°00’-48°10’N), Urumqi, Xinjiang, China and Fuhai County, Altay, Xinjiang, China. See Supplementary Table 8 for sample information.

**Herdsmen’s saliva, feces, height and weight**: The current study was approved by Ethics Committee of People’s Hospital of Xinjiang (KY2022022302) and conducted in accordance with the Health Research Authority guidelines. Collected from Fuhai County, Altay Region, Xinjiang, China. See Supplementary Table 9-10 for sample information.

### Experiment of rats

#### Preparation of camel whey and Bovine whey

Centrifuged fresh camel milk and bovine milk at 5000 r/min for 20 min, discarded fat and precipitate, and took the middle layer to obtain skim milk. Collected skim milk from each centrifuge tube, heated it in 40% water bath for 20 min, then adjusted the pH to 4.6 with 10% glacial acetic acid, and placed it in a refrigerator at 4℃ overnight. The overnight camel skim milk and bovine skim milk were placed in a centrifuge tube, centrifuged at 8 000 r/min for 20 minutes, and the intermediate whey was collected twice. Poured the centrifugal camel whey and bovine whey into a culture dish, marked and sealed it, froze it at -80℃ for 12 hours. Took out the culture dish filled with camel whey and bovine whey the next day, punched the hole on the packed culture dish with a sterile toothpick, put it into a pre-heated freeze dryer for drying, and finally collected the freeze-dried powder of camel whey and bovine whey.

#### Modeling and grouping of rats

The current study was approved by the Ethics Committee of the First Affiliated Hospital of Xinjiang Medical University (IACUC-20200620-01). SD rats were randomly divided into normal control group, diabetic model group, camel whey prevention group and bovine whey prevention group. The rats in the normal control group were fed normally, and the rats in the other groups were fed with 45% high-fat diet. At the same time, the camel whey prevention group and the bovine whey prevention group were intragastrically fed with 200 mg camel whey freeze-dried powder and bovine whey freeze-dried powder every day for 6 weeks, and the model was established after 6 weeks. Rats in diabetes model group, bovine whey prevention group and camel whey prevention group were weighed first and then injected with 35mg STZ. After 7 days of modeling, the fasting blood glucose value was detected by blood glucose meter, and the rats whose fasting blood glucose value was greater than 11.1mmol/dL were used in the follow-up experiment, See Supplementary Table 11 for sample information.

The diabetic rats were divided into diabetic model group and positive drug group according to their body weight and blood glucose concentration. According to the dose of camel whey and bovine whey, the diabetic model group was again divided into high-dose camel whey group, low-dose camel whey group, high-dose bovine whey group and low-dose bovine whey group. The rats that meet the criteria were singled out to continue the follow-up experiment. There were 9 groups: normal control group, diabetic model group, positive drug group, camel whey prevention group, camel whey low-dose group, camel whey high-dose group, bovine whey prevention group, bovine whey low-dose group, and bovine whey high-dose group. Rats in positive drug group were given 200 mg hydrochloric acid metformin. Camel whey prevention group, camel whey high-dose group and bovine whey prevention group, high-dose bovine whey group were still intragastrically infused with 200 mg camel whey and bovine whey freeze dried powder. Camel whey low-dose group and bovine whey low-dose group were given intragastric administration of 50 mg camel whey and bovine whey freeze dried powder. Normal and diabetic model rats were intragastrically infused with normal saline. All the above treatments were once a day for 6 weeks. During this period, the fasting blood glucose and body weight of rats were measured and recorded every week.

#### Anatomy and sampling of rats

The SD rats which reached the standard after grouping were killed by the method of cervical vertebra removal. Used anatomical scissors to cut the end of the rat’s abdomen along the midline of the abdomen and removed the intestinal part from the lower end of the stomach to the anus. The intestinal tract of rats was divided into four parts: duodenum, ileum, cecum and colon. Then the contents of various parts of rat intestine were extruded into 15ml aseptic EP tube by hand extrusion, marked and stored at-80 ℃. See Supplementary Table 12 for sample information.

#### Collection of rat blood samples

Before the standard SD rats were killed, the blood of normal control group, high dose camel whey group, positive drug group and diabetic model group were collected by retroorbital venous plexus method, marked and stored at -80℃. See Supplementary Table 13 for sample information.

After all the above samples were collected, the samples of plants, camel digestive tract, camel milk, camel feces, rat intestinal tissue and herdsman saliva and feces were entrusted to Meiji Biomedical Technology Co., Ltd. Based on the third-generation sequencing platform of Ilumina, 16s high-throughput sequencing and macrogenomic sequencing were carried out to explore the information of bacterial community composition, abundance, similarity and difference, flora function and so on. The blood samples of rats were entrusted to Zhongke Xinsheng Biotechnology Co., Ltd. for serum non-targeted metabonomic sequencing. The composition and changes of serum metabolites of diabetic rats under different treatments were analyzed. And it was combined with microbiology to explore the interaction between camel milk and gut microbes and type 2 diabetic rats.

### Data processing and analysis

16s high-throughput sequencing results were used Uparse (v.7.1) to cluster classification operation units (Operational taxonomic units, OTU) at 97% similarity to get the representative sequence of OTU. Then Alpha diversity analysis was carried out by Mothur (v.1.30.2) to reflect species richness and diversity in the samples. Beta diversity analysis was carried out by Qiime (v.1.9.1) to compare the community composition of the tested samples, and R language (version3.3.1) was used to analyze the species composition of the tested samples.

In the results of metagenome sequencing, the original sequencing data were controlled by fastp software, and the short segment sequences obtained by quality control were assembled by Multiple_Megahit. The assembly results were clustered by CD-HIT software to construct non-redundant gene set, and the high-quality reads of each sample were compared with non-redundant gene set using SOAPaligner software (default parameter: 95% identity), and the abundance information of genes in the corresponding samples was calculated. Then the non-redundant gene set sequence was compared with KEGG gene database (GENES) by DIAMOND (parameter: blastp; E-value≤ 1e-5). According to the gene abundance sum of KO, Pathway, EC and Module, the abundance of this functional category was calculated, and based on the corresponding abundance data table, the functional composition of microbes in the tested samples was analyzed by R language.

The original data of serum metabolic sequencing was converted into mzXML format by ProteoWizard, and then peaked alignment, retention time correction and peaked area extraction were performed by XCMS program. The structure of metabolites was identified by accurate mass number matching (<25ppm) and secondary spectrum matching to search the database (56). For the data extracted by XCMS, deleted the ion peaks with missing values > 50% in the group. The software SIMCA-P14.1 (Umetrics,Umea,Sweden) was used for pattern recognition. After the data was preprocessed by Pareto-scaling, the dimension of the multivariable original data was reduced by PCA (principal component analysis). The grouping trend (intra-group and inter-group similarity and difference) and outliers (whether there are abnormal samples) of the observed variables in the data set were analyzed. The differential metabolites were screened by the multiple of difference (Fold change) and T test (Student’st-test) obtained by univariate analysis.

## Author’s contributions

H. Y. participated in the design, performance, and analysis of most studies and the drafting of the manuscript. J. Z. and R. W. participated in the design and implementation of microbial studies and analysis, and the drafting of the manuscript. L. Z., Y. K., R. L., and Z. Y. assisted in all rat studies. J. Y. and Y. L. performed all mass spectrometry analyses. Y. Q., X. L., and X. W. assisted with statistical analyses. P. Y. and J. W. performed gut microbial composition analyses. X. X., L. X., and H. N. provided input on the experimental design and were involved in thoughtful discussions. G. C., T. S., and F. G. conceived the project idea and participated in the design of experiments, Sample Collection. All authors critically reviewed and edited the manuscript.

## Acknowledgements

This research was supported by the National Key Research and Development Program of China (grant 2019YFC1606102), the Outstanding Young Scientific and Technological Talents Training Program of Xinjiang Autonomous Region (grant 2020Q002), National Natural Science Foundation of China (grant U2003305, 31860018).

## Ethics approval and consent to participate

For human research, it was approved by the Ethics Committee of Xinjiang People’s Hospital (KY2022022302) and conducted according to the guidelines of health research institutions. The use of experimental animals was approved by the Ethics Committee of the First Affiliated Hospital of Xinjiang Medical University (IAUC-20200620-01).

## Data availability

All data needed to evaluate the conclusions in the paper are present in the paper and/or the Supplementary Materials. The data that support the findings of this study are available as Mendeley Data, V1, https://doi.org/10.17632/4w8n8n96tc.1.

